# Importance of genetic architecture in marker selection decisions for genomic prediction

**DOI:** 10.1101/2023.02.28.530521

**Authors:** Rafael Della Coletta, Samuel B. Fernandes, Patrick J. Monnahan, Mark A. Mikel, Martin O. Bohn, Alexander E. Lipka, Candice N. Hirsch

## Abstract

Breeders commonly use genetic markers to predict the performance of untested individuals as a way to improve the efficiency of breeding programs. These genomic prediction models have almost exclusively used single nucleotide polymorphisms (SNPs) as their source of genetic information, even though other types of markers exist, such as structural variants (SVs). Given that SVs are associated with environmental adaptation and not all of them are in linkage disequilibrium to SNPs, SVs have the potential to bring additional information to multi-environment prediction models that are not captured by SNPs alone. Here, we evaluated different marker types (SNPs and/or SVs) on prediction accuracy across a range of genetic architectures for simulated traits across multiple environments. Our results show that SVs can improve prediction accuracy by up to 19%, but it is highly dependent on the genetic architecture of the trait. Differences in prediction accuracy across marker types were more pronounced for traits with high heritability, high number of QTLs, and SVs as causative variants. In these scenarios, using SV markers resulted in better prediction accuracies than SNP markers, especially when predicting untested genotypes across environments, likely due to more predictors being in linkage disequilibrium with causative variants. The simulations revealed little impact of different effect sizes between SNPs and SVs as causative variants on prediction accuracy. This study demonstrates the importance of knowing the genetic architecture of a trait in deciding what markers and marker types to use in large scale genomic prediction modeling in a breeding program.

**Key message:** We demonstrate potential for improved multi-environment genomic prediction accuracy using structural variant markers. However, the degree of observed improvement is highly dependent on the genetic architecture of the trait.

## Introduction

Breeding and releasing a new plant variety can take years or even decades (Challinor et al. 2016; Voss-Fels et al. 2019). One important tool available for breeders to increase the efficiency of a breeding program is genomic prediction, where the performance of individual plants is predicted based solely on their genetic information (Meuwissen et al. 2001). It allows plant breeders to discard unwanted genotypes early in the breeding process, reduce breeding cycle time, and more efficiently allocate resources in their breeding program (Lorenz et al. 2011). Genomic prediction has been successfully implemented in many commercial breeding programs mainly because genotyping on a large scale is less expensive than phenotyping. However, prediction accuracy is variable, highly dependent on the population structure, diversity, and traits being predicted, and these challenges escalate when predicting traits across multiple environments (Burgueño et al. 2012; Combs and Bernardo 2013; Lian et al. 2014). These issues have motivated many research groups to study new ways to capture and use more relevant information in genomic prediction models.

Prediction models used by breeding programs rely heavily on single nucleotide polymorphisms (SNPs) mainly because they are abundant in the genome and easy to measure via low-cost molecular assays. Studies in maize and rice showed that adding transcriptomic (mRNAs and small RNAs) and metabolomic data improved prediction accuracy, but it was highly dependent on the trait being evaluated (Xu et al. 2016; Guo et al. 2016; Westhues et al. 2017; Schrag et al. 2018). For example, Schrag et al. (2018) observed that using only transcriptomic markers improved prediction accuracy by up to 14% compared to only genomic markers when predicting grain yield in maize hybrids, but accuracy was approximately 10% lower when predicting grain dry matter content. Another study showed that genomic and transcriptomic markers have similar performance, but when using only the most informative marker from each type, prediction accuracy increased by about 6% (Azodi et al. 2020). More recently, it was shown that using imputed gene expression via haplotype associated RNA expression (HARE) in genomic prediction models is more stable and more accurate than measured gene expression (Giri et al. 2021). Although the authors showed that HARE predictions have similar accuracy to SNP predictors, HARE may be particularly useful for cross-population predictions since they do not require that all SNPs are shared across populations. All of these studies highlight that although SNPs are able to predict traits with relatively decent accuracy, maximizing prediction accuracy will likely require the use of additional marker types.

Another type of marker that has been largely ignored in prediction models are structural variants (i.e., deletions, insertions, duplications, inversions, translocations; hereafter collectively called SVs). Initial analysis with structural variants identified from 787 yeast genomes revealed that genomic prediction with pan-genomic open reading frames (ORFs) was, on average, two times more accurate than SNPs across 35 different traits (Li and Simianer 2020). Advances in sequencing technologies and bioinformatics tools are allowing a more accurate identification of these variants (Ho et al. 2019), which are relatively common in crop species (Montenegro et al. 2017; Gao et al. 2019; Liu et al. 2020, 2022; Hufford et al. 2021; Varshney et al. 2021; Rijzaani et al. 2022; Shang et al. 2022). In addition, many studies have shown that SVs are associated with tolerance to biotic (Cook et al. 2012; Zuo et al. 2015) and abiotic stresses (Sutton et al. 2007; Knox et al. 2010; Maron et al. 2013), changes in flowering time (Nitcher et al. 2013; Würschum et al. 2015), and plant architecture (Zhou et al. 2009; Studer et al. 2011). Importantly, it has been demonstrated that SVs can explain phenotypic variation that SNPs cannot (Song et al. 2020; Hufford et al. 2021), and that not all SVs are in linkage disequilibrium with a SNP that effectively “tags” the SV (Chia et al. 2012; Stuart et al. 2016; Yang et al. 2019; Qiu et al. 2021). These findings raise the question of whether SVs can provide additional genetic information to prediction models not captured by SNPs. One study in maize attempted to model copy number variants (CNVs) in genomic prediction models with maize hybrids and observed an increase of approximately 6% in accuracy when predicting plant height under low nitrogen conditions with SNPs and CNVs (Lyra et al. 2018). More recently, Chen et al. (2021) also tested SVs in genomic prediction models in cattle, and concluded that there was no benefit in using such markers for predicting dairy production traits. Although important, the above studies with empirical data do not fully investigate the potential of SVs in genomic prediction models as only a few traits with unknown genetic architecture were analyzed, which may limit the generalization of their results.

In this study, we performed a comprehensive evaluation of the usefulness of structural variants in multi-environment genomic prediction models using a simulation approach to guide the interpretation of results from future empirical data. First, we simulated phenotypes using real genotypic information from maize recombinant inbred lines, varying heritability, number of quantitative trait loci (QTLs), type of causative variant (SNPs and SVs), and variant effect sizes. Then, we tested different types of markers (SNPs and/or SVs) as predictors of genomic prediction models and evaluated their accuracy for each simulated genetic architecture. The overall goal of this study was to determine if there are genetic architectures for which the use of higher-cost SV markers would improve prediction accuracy enough to justify their inclusion in large scale genetic prediction modeling in a breeding program.

## Materials and methods

### Genotypic data

To simulate traits with real-world population structure, we used genotypic information from maize lines that have been extensively used in commercial breeding programs and are representative of their diversity for trait simulations.The population of 333 F7 RILs, which has been previously described (Della Coletta et al. 2023), was generated from half diallel crosses of six maize inbred lines including: B73, PHG39, PHG47, PH207, PHG35, and LH82. The parental lines and the 333 F7 RILs were previously genotyped with a custom Illumina Infinium 20K SNP chip (see ‘Availability of data and material’ section). The parental lines have also been previously genotyped using RNA-seq reads for over 20 million SNPs (Mazaheri et al. 2019).

In addition to these existing SNP variant calls, structural variants were identified in the parental lines from whole genome sequencing data available in Qiu et al. (2021). Adapter sequences were trimmed using CutAdapt v1.18 with default parameters (Martin 2011), and low quality bases were removed using sickle v1.33 with default parameters (Joshi and Fass 2011). Reads were mapped to the ten assembled chromosomes in the B73 v4 reference genome (Jiao et al. 2017) using Speedseq v0.1.2 (Chiang et al. 2015), a parallelized version of bwa (Li and Durbin 2009; Li 2013) that simultaneously isolates the split reads and discordant paired reads with unusually large insert sizes. To be included in the split reads, we required a minimum of 20 non-overlapping base pairs between two alignments and allowed at most two alignments for a particular read. Duplicate reads were removed during the mapping process.

To identify and genotype structural variants, we utilized Lumpy v0.2.13 (Layer et al. 2014) as implemented by the ‘speedseq sv’ command. The analysis of SVs was restricted to the gene space (± 2kb on gene boundaries), using bed files containing regions to be excluded (via the ‘-x flag’). Lumpy was initially run on each sample individually and estimated copy number via Lumpy’s internal calls to CNVnator (Abyzov et al. 2011) using a window size of 300bp. The SVtools v0.5.1 pipeline (Larson et al. 2019) was used to merge individual VCFs, re-genotype all samples at all sites, and prune redundant variants. SVs were required to be within 500bp of one another in order to remain in a cluster. The final step in the SVtools pipeline is to re-classify low quality SVs as BND (the generic ‘breakend’ annotation) as well as SVs that overlap repetitive elements as MEI (‘Mobile Element Insertion’), which was done using the svtools classify subroutine with the ‘large_sample’ option. All BND variants were removed from the final variant file and MEI variants were recorded with the original DEL annotation (Supplemental File 1). For compatibility with available bioinformatics tools and genomic prediction models, SV information was reduced to a binary state (i.e., presence/absence of SV), and the middle position of the SV was used as its point position in the genome. Information regarding SV type and breakpoint positions were stored in the final marker names.

### Projection of whole genome variants to RILs

The ∼20 million SNPs and ∼10,000 SVs from the deep parental information were projected onto the 333 RILs using the 20,000 SNP chip markers to define haplotype blocks. Prior to projections, parental SNPs from RNA-seq that were identified within the boundaries of deletions of 100kb or less and monomorphic markers were removed. The genotypic information for each diallel family was then split apart. SNP chip markers with segregation distortion within families (Chi-square test, FDR-corrected p-value < 0.05) were identified using R/qtl v1.48 (Broman et al. 2003) and removed. Additionally, within families SNPs that were within a parental deletion of 100kb or less (99% of all deletions) were removed. To correct for errors during genotyping, a sliding window approach was implemented as described by Huang et al. (2009) with a 15-bp window, 1-bp step size, and a minimum of five markers per window.

Projections were done on each diallel family separately using the FILLIN plugin from TASSEL v5.2.56 (Bradbury et al. 2007) as described in Hufford et al. (2021). Briefly, parental haplotypes were created with FILLINFindHaplotypesPlugin (-hapSize 200000 -minTaxa 1) and then projected onto missing genotypes in the RILs with FILLINImputationPlugin (-hapSize 200000 -hybNN false). Finally, we re-applied the previously described sliding window approach (Huang et al. 2009) with a 45-bp window slide, 1bp step size, and a minimum of 15 markers per window. The projection results of all diallel families were combined into a single dataset, and monomorphic markers across all RILs were removed (Supplemental File 2).

### Linkage Disequilibrium structure of RIL population

Linkage Disequilibrium (LD) was calculated using plink v1.90b6.16 (Purcell et al. 2007) as previously described (Qiu et al. 2021) for markers with less than 25% missing data. When calculating LD between SNPs and SVs, the window size was limited to 1kb to reflect the LD decay in this population. The reason for this was to find SNPs that were close enough to SVs to avoid LD due to population structure, but far enough to allow recombination between SNPs and SVs.

### Environmental data

Weather data from five locations in the U.S. Midwest (Iowa City, IA, Bloomington, IL, Champaign, IL, Janesville, WI, and Saint Paul, MN) from April 2020 to October 2020 were obtained using the R package EnvRtype v1.0 (Costa-Neto et al. 2021) (Table S1). A weather covariable matrix with 17 weather parameters (Table S2) in 10-day intervals through the growing season was obtained and transformed into a variance-covariance kinship matrix for environmental relatedness with the env_kernel function in EnvRtype v1.0 (Costa-Neto et al. 2021), and further converted to a correlation matrix (Table S3) with cov2cor function in R v3.6 (R Core Team 2019).

### Trait simulation across multiple environments

The R package simplePHENOTYPES v1.3 (Fernandes and Lipka 2020) was used to simulate traits across five different environments for the RIL population described above (Supplemental File 3). This package was originally designed for simulating pleiotropic traits based on a user-defined genetic architecture. However, simulating multiple traits for an individual is equivalent to simulating the performance of an individual across multiple environments when the loci controlling trait variation are kept the same. Multi-environment traits were simulated with three replicates for each inbred line (additive effects only) with different heritabilities (0.3 or 0.7), different number of loci controlling trait variation (10 or 100), and different causative variant types (SNPs only, SVs only, or both) (Fig. 1a-b).

**Fig. 1.**
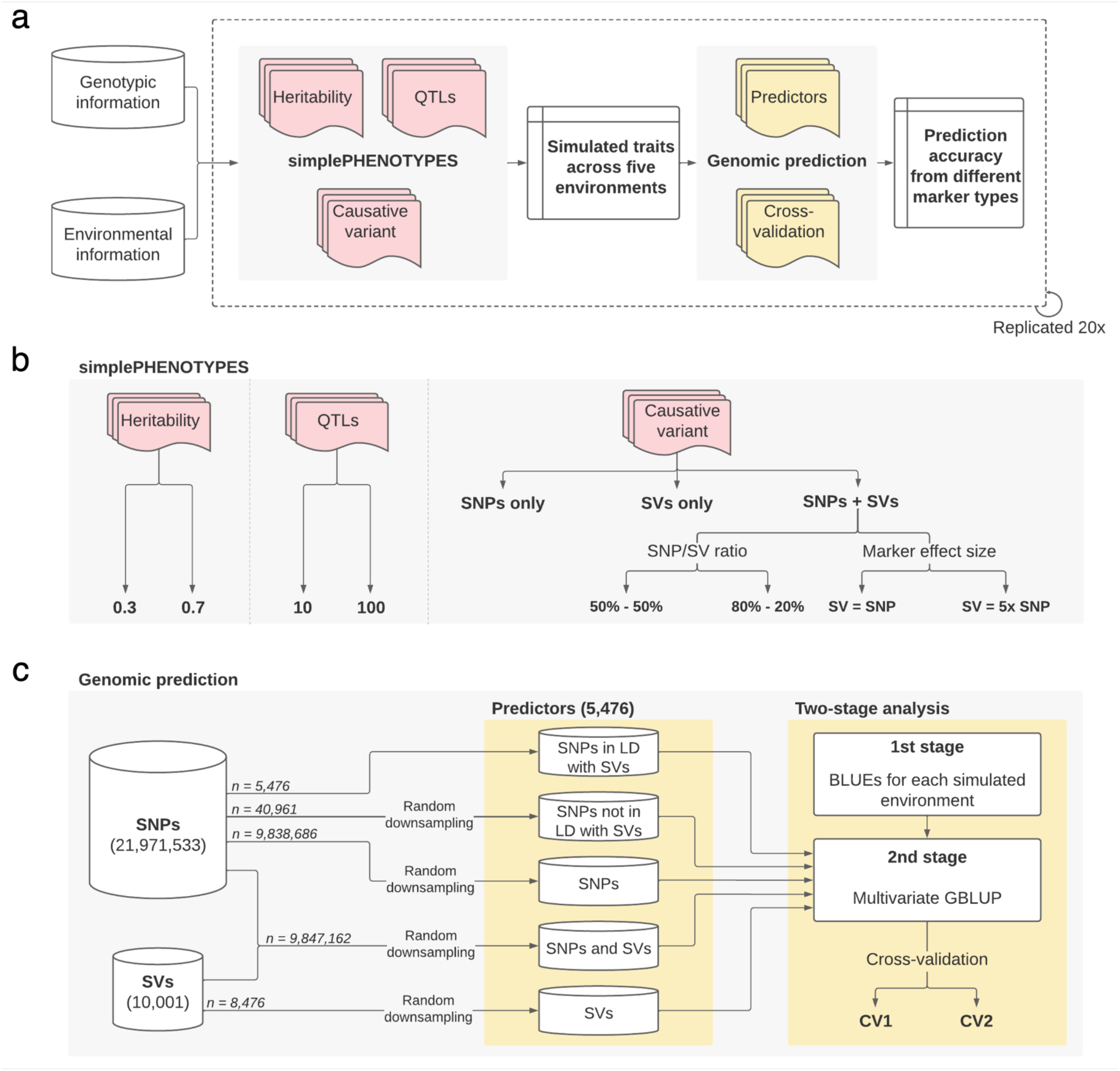
Methodology overview for simulating traits and testing performance of different marker types in genomic prediction models. **a** Genotypic data from maize inbred lines and environmental correlation matrix were loaded into the R package simplePHENOTYPES for trait simulation. Genomic prediction models were run with the simulated data and prediction accuracy was evaluated for different marker types with distinct cross-validation schemes. The whole process was repeated 20 times; **b** In simplePHENOTYPES, traits were simulated with a factorial combination of different heritabilities (0.3 or 0.7), QTL number (10 or 100) and causative variants (SNPs and/or SVs) across multiple environments with genotype-by-environment interactions; **c** For genomic prediction, five different marker types were used: SNPs only, SVs only, SNPs and SVs, SNPs in LD to SVs, and SNPs not in LD to SVs. These predictors were randomly subsampled from the total amount of projected markers to keep prediction comparisons fair. Finally, prediction accuracy of each marker type for each simulated trait was obtained using a two-stage genomic prediction model with two cross-validation schemes (CV1 and CV2)

Environmental information was incorporated as a residual correlation matrix in simplePHENOTYPES, and each QTL was given the same small additive effect on phenotypic variation across all environments. Genotype-by-environment interactions (GxE) was simulated by adding random effects (drawn from a (0, 0.3) distribution and bound between -1 and 1) to each QTL at each environment. The following conditions were also imposed when adding GxE effects: (1) 10% of QTLs were allowed to change the signs of effects across environments, (2) 10% of QTLs had no effect at all in a particular environment, and (3) 10% of QTLs had constant effects across environments (i.e., no GxE). The 10% of QTL for which these conditions were applied were different for each of the conditions. To confirm that GxE was successfully simulated in each population, analysis of variance (ANOVA) was performed with the R package lme4 (Bates et al. 2015) using the linear model:

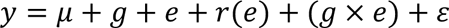

where 𝑦 is the simulated trait values, 𝜇 is the overall mean, 𝑔 is the fixed genotypic effect of an inbred line, 𝑒 is the random environmental effect, 𝑟(𝑒) is the random replicate effect within environment, 𝑔 × 𝑒 is the random effect of genotype-by-environment interactions, and 𝜀 is the random error term (Supplemental File 4).

When both SNPs and SVs were selected as causative variants (SNP-to-SV ratio of 1:1 or 4:1), we simulated scenarios where SVs had the same effect size as SNPs or 5x larger effect size compared to SNPs. We added such scenarios to test the hypothesis that SVs may have a bigger impact on phenotypes than SNPs.

For all 24 scenarios (i.e., the combination of heritability, number of loci, causative variant type, and SNP-to-SV ratio when applicable; Table S4), we performed 20 independent simulation replicates with different QTLs selected each time with replacement (Fig. 1a-b) to reduce any bias introduced by chance when selecting the QTLs that control a trait.

### Genomic prediction models

Genomic prediction of each of the 24 simulated populations described above was performed using (1) only SNP markers, (2) only SV markers, (3) all SNP and SV markers, (4) only SNP markers in LD with an SV, and (5) only SNP markers not in LD with an SV (Fig. 1c). The last two types of markers were included to test the ability of certain SNPs being used as proxies for SVs. A SNP was considered in LD with an SV if r^2^ > 0.9. Markers that were selected as causal variants during trait simulations were not masked and therefore could be selected as predictors. Due to the difference in the absolute numbers of marker types (millions of SNPs vs thousands of SVs), we randomly downsampled the same number of predictors according to the predictor type with the lowest number of markers (n = 5,476) and repeated this process of selecting markers 20 times to reduce bias due to random chance (Supplemental File 5). This totaled 4,800 genomic prediction models (24 scenarios x 5 types of predictors x 2 cross-validations x 20 replications) that were run (Fig. 1c)

Genomic prediction models were run in two stages using ASReml-R v4.1 (Butler et al. 2017) as described by Fernandes et al. (2018). In the first stage, we obtained best linear unbiased estimates (BLUEs) for each environment to account for GxE effects with the following model:

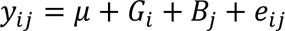

where 𝑦_𝑖𝑗_ is the simulated trait value of genotype 𝑖 in replicate 𝑗, 𝜇 is a constant, 𝐺_𝑖_ is the fixed effect of the 𝑖th genotype, 𝐵_𝑗_ is the random effect of the 𝑗th replicate with 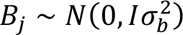, 𝑒_𝑖𝑗_ is random effect of residuals with 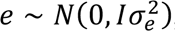, and 𝐼 is an identity matrix.

In the second stage, we ran multivariate genomic best linear unbiased prediction (GBLUP) models to evaluate the performance of different marker types with the following model:

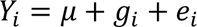

where 𝑌_𝑖_ is the vector of multivariate responses associated with genotype 𝑖 (𝑖 = 1,2, …, 𝑛) in which 𝑌_𝑖_ = [𝑌_𝑖1_, 𝑌_𝑖21_, …, 𝑌_𝑖𝑝_]^т^, 𝜇 is the vector of the constants associated with each environment with 𝜇 = [𝜇_1_, 𝜇_2_, . . ., 𝜇_𝑝_]^т^, 𝑔_𝑖_ is the vector of random effects of genotype 𝑖 associated with each environment in which 𝑔 = [𝑔_1_, 𝑔_2_, …, 𝑔_𝑖_, …, 𝑔_𝑛_]^т^, 𝑔 ∼ 𝑁_𝑛𝑝_(0, 𝐺 ⊗ 𝐴) with 𝐺 as the variance-covariance matrix for genetic effects, ⊗ as the Kronecker product and 𝐴 as the realized additive relationship matrix calculated from the genotypic data using the Endelman and Jannink method (Endelman and Jannink 2012) implemented in the A.mat function from the R package rrBLUP v4.6.1 (Endelman 2011), 𝑒_𝑖_ is the vector of random effects of residuals in which 𝑒 = [𝑒_1_, 𝑒_2_, …, 𝑒_𝑖_, …, 𝑒_𝑛_]^т^, 𝑒 ∼ 𝑁_𝑛𝑝_(0, 𝐼 ⊗ 𝑅) with 𝑅 as the variance-covariance matrix for residual effects, and 𝐼 as an identity matrix. Both SNPs and SVs were considered biallelic for the calculation of the additive relationship matrix and were coded as {– 1,0,1}. For SNPs, -1 represents one of the homozygous states, 0 represents the heterozygous state, and 1 represents the alternative homozygous state. Likewise, for SVs -1 represents one of the homozygous states (e.g. presence or absence of the SV), 0 represents the heterozygous state, and 1 represents the alternative homozygous state. The 𝐺 matrix was assumed to have a diagonal structure (i.e., genotypes were allowed to have different variance across environments but without co-variance between environments) while the 𝑅 matrix was assumed to be unstructured to take advantage of correlations among environments. This strategy was chosen as a compromise between computational efficiency and the amount of information to be used in each of the predictions.

Prediction accuracy was calculated using two different 5-fold cross validation schemes: CV2 and CV1 (Burgueño et al. 2012). CV2 was used to determine how well genotypes with phenotypic information in some environments are predicted, whereas CV1 evaluated how accurately we predicted completely untested genotypes. In both schemes, we randomly divided our simulated dataset into five different groups and predicted the phenotypes of one group based on the data from the other four. After the phenotypes of all five groups were predicted, we determined prediction accuracy by calculating the average Pearson’s correlation between observed and predicted values. The cross-validation process was repeated three times to reduce bias due to random chance when assigning genotypes to one of the five groups (Fig. 1c).

### LD and prediction accuracy

We also sought to evaluate the extent to which LD between QTL and predictors impacts prediction accuracy. To this end, we calculated LD between all QTLs and all predictors as described above but without a window size limit, and plotted the distribution of LD against prediction accuracy. Given the total number of markers used in genomic prediction models, we expected that most of the QTLs would be in high LD with at least one marker, which would reduce the ability to detect meaningful signals between LD and prediction accuracy. Thus, we designed another simulation experiment to better answer this question. For this purpose, we randomly downsampled the total number of markers to approximately 25% to reduce computational resources while making sure the minimum allele frequency distribution of SNPs and SVs had a similar pattern. Then, we calculated LD among all remaining markers in 5Mb windows as described above. Finally, we simulated new traits (100 QTLs; 0.3 or 0.7 heritability; SNPs or SVs as causative variants) across the same five environments described above using the markers that had LD information. The simulation was repeated three times, where each time a different set of markers was selected as the causative variant (Supplemental File 6). From the same pool of markers with LD information, we selected 500 markers that were in low LD (r^2^ < 0.5), moderate LD (0.5 ≤ r^2^ < 0.9), or high LD (r^2^ ≥ 0.9) to a QTL and ran genomic prediction models as described above (Fig. S1). To avoid model convergence problems due to highly correlated markers in the moderate and high LD categories, we selected only one marker with the highest LD with a QTL (i.e., 0.5 < r^2^ < 0.9 for moderate LD category, and r^2^ > 0.9 for high LD category) and randomly selected the remaining markers from markers in the lower LD categories (i.e., r^2^ < 0.5 for the moderate LD category, and r^2^ < 0.9 for the high LD category). This experiment was repeated 10 times (Supplemental File 7).

## Results

### Not all SVs are in high linkage disequilibrium to a SNP

To maximize the amount of genetic information available in the RILs in a resource efficient way, we projected markers from whole genome resequencing of the parents of the diallel families to their respective RILs. Approximately 93.6% of all markers were successfully projected among all families with an average accuracy of 94.6% (Fig. S2), resulting in a total of 21,971,533 SNPs and 10,001 SVs (9,540 deletions, 241 inversions, and 220 duplications) in the RIL population. The presence of SVs near centromeric regions was depleted compared to SNPs, which had a more uniform distribution along the chromosomes (Fig. S3).

The usefulness of SVs in genomic prediction models is highly dependent on the LD structure of the population in which the predictions are made. For example, if all SVs are in high LD to SNPs, then adding SVs in genomic prediction will add little information to the models because all phenotypic variation associated with those SVs would have already been captured by SNPs. We investigated the LD structure in this RIL population and found that LD decays very rapidly. While this is a RIL population with limited rounds of recombination, due to the multiparent design, LD declines more rapidly with the median r^2^ values between SNPs within 400bp from each other already lower than 0.5 (Fig. 2a). LD analysis between SNPs and SVs revealed approximately 24% of SVs were not tagged by a SNP (r^2^ < 0.9; Fig. 2b). Consequently, for this population, SVs could bring additional information into genomic prediction models.

**Fig. 2.**
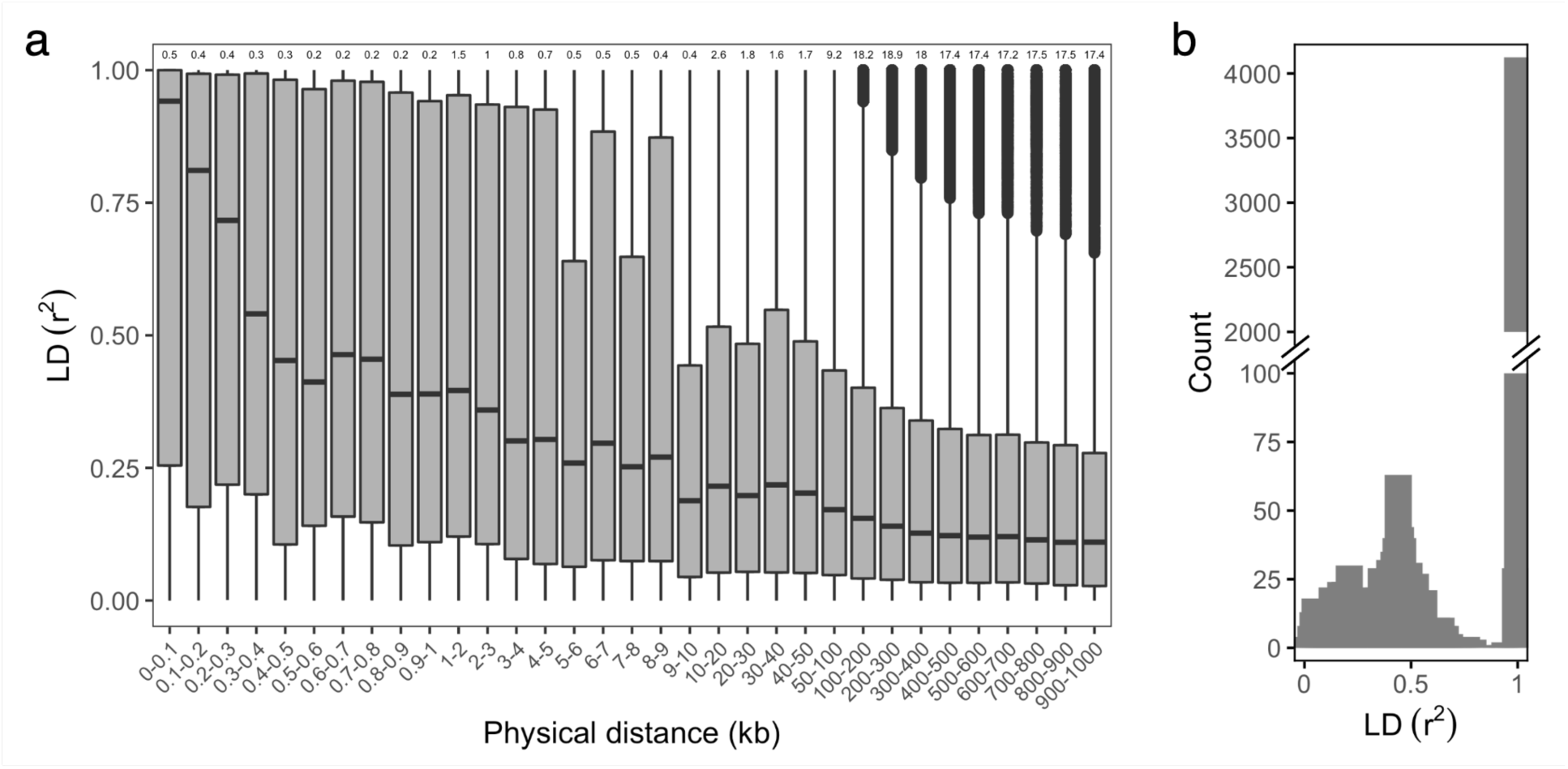
Linkage disequilibrium (LD) structure in a maize population with 333 recombinant inbred lines derived from diallel crosses of six parental lines. **a** Boxplot with pairwise r^2^ values between SNP chip markers within 1Mb from each other. Numbers above each box represent the number (in thousands) of pairwise LD values of markers within a certain physical distance; **b** Distribution of r^2^ values between SNPs and SVs for the SNP in highest LD to each SV

### SVs can improve genomic prediction accuracy, but it depends on the genetic architecture of the trait

To test if SVs bring additional information into genomic prediction models, we conducted genomic prediction on eight simulated populations with varying genetic architectures controlled by either SNPs or SVs, and with different marker types as predictors (i.e.. SNPs only, SVs only, both SNPs and SVs, SNPs in LD with SVs, and SNPs not in LD with SVs) (Fig. 3a, Table S5, Supplemental File 8). As expected, higher trait heritability and higher number of QTLs were associated with higher prediction accuracy across all of the predictor types. On the extremes, the mean accuracy across different predictor types for simulated traits with 10 QTLs at *h^2^* = 0.3 was ∼55.7%, while when the trait architecture was 100 QTLs at *h^2^* = 0.7 the average accuracy across the different predictor types improved to ∼79.4%.

**Fig. 3.**
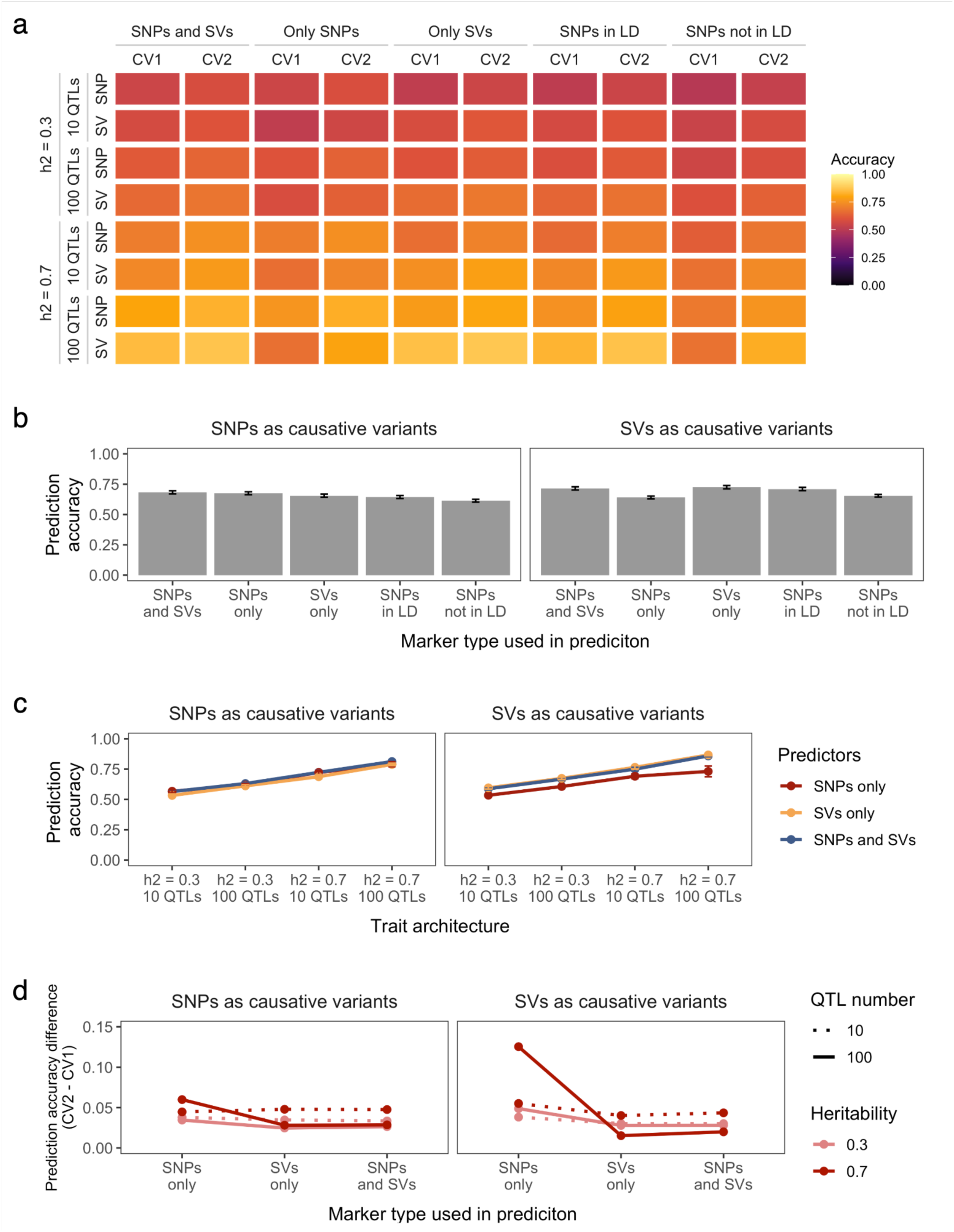
Genomic prediction accuracy of different marker types (SNPs and SVs, SNPs only, SVs only, SNPs in LD to SVs, or SNPs not in LD to SVs) from two cross-validation schemes (CV1 and CV2) for simulated traits where either SNPs or SVs were the causative variants. **a** Heatmap summarizing the average performance of each predictor type from two cross-validation schemes (x-axis) for each simulated genetic architecture with different heritabilities and number of QTLs controlling trait variability (y-axis); **b** Average prediction accuracy of marker types across all genetic architectures; **c** Average prediction accuracy of marker types for different genetic architectures; **d** Average prediction difference between the two cross-validation schemes of marker types for different genetic architectures

Overall, when only SNPs control trait variation, using SNP predictors resulted in slightly better prediction accuracy than SV predictors (Fig. 3b, Table S5), with the largest difference observed when a trait had lower heritability and was controlled by fewer variants (Table S5). In this case, the difference in average prediction accuracy between SNP predictors and SV predictors was 3.5% in CV2 (Welch’s t-test, t(36.257) = 2.485, p-value = 0.017) and 3.3% in CV1 (Welch’s t-test, t(37.21) = 1.875, p-value = 0.0687). As both heritability and number of QTLs increased, including both SNPs and SVs or only SVs as predictors resulted in more similar accuracies as with SNPs alone (Fig. 3c). In contrast, when a trait was controlled by SVs exclusively, using SVs as predictors (either by themselves or in conjunction with SNPs) resulted in higher accuracy than using only SNPs in all scenarios (Fig. 3c, Table S5). In this case, the difference in average prediction accuracy between SNP predictors and SV predictors was as high as 8% in CV2 (Welch’s t-test, t(19.813) = 7.566, p-value < 0.001) and 19% in CV1 (Welch’s t-test, t(16.755) = 6.674, p-value < 0.001) for highly heritable traits controlled by many QTLs.

We also tested the ability of SNP markers in LD to SVs (r^2^ > 0.9) to be used as a proxy of SVs in genomic prediction models, and we found that this marker type resulted in a very similar overall accuracy pattern of SVs predictors, although with slightly lower values (Fig. 3b, Table S5). The average difference between SVs as predictors and SNPs in LD to SVs as predictors for traits controlled solely by SNPs or SVs were ∼ 1.8% for CV2 (Welch’s t-test, t(307.97) = 1.513, p-value = 0.131) and ∼ 2.3% for CV1 (Welch’s t-test, t(307.82) = 1.877, p-value = 0.061). On the other hand, SNPs that were not in LD to any SVs (r^2^ < 0.9) were the worst predictors when SNPs were the causative variants of traits, but performed similarly to general SNP predictors when SVs controlled trait variation (Fig. 3b, Table S5).

It is expected that results from CV2 are better than CV1 in all scenarios due to the latter having to predict the performance of genotypes with no phenotypic information in any environment. Interestingly, using SVs as predictors when SVs were the causative variants reduced the difference between CV1 and CV2 in terms of accuracy, especially at higher heritability and QTL numbers (Fig. 3d, Table S5). For instance, when a trait was controlled by 100 SVs at *h^2^* = 0.7, the average difference was approximately 1.5%, whereas the average difference between CV1 and CV2 in the same scenario but when only SNPs were used for predictions was approximately 12.5% (Welch’s t-test, t(15.162) = 7.386, p-value < 0.001).

### Adding SVs to prediction models is generally beneficial when trait genetic architecture is controlled by both SNPs and SVs

Given that it is unlikely, especially for traits controlled by multiple QTLs, that only SNPs or SVs will be the causative variants, we also tested 16 simulated scenarios where both SNPs and SVs control trait variation (Fig. S4; Table S6; Supplemental File 10). When half of the causative variants are SNPs and the other half are SVs and both marker types have the same effect size, there is nearly no difference between using SVs only or SNPs only as predictors in the case when there are few QTLs and the heritability is low (accuracy difference of ∼ 1.1% for CV2 (Welch’s t-test, t(33.527) = 1.007, p-value = 0.321 and ∼ 1.6% for CV1 (Welch’s t-test, t(33.675) = 1.322, p-value = 0.195); Fig. 4a, Table S6, Fig. S4). However, similar to scenarios where only SNPs or SVs control trait variation, using SVs in prediction models results in better accuracy than SNP predictors as the number of QTLs and heritability increase (accuracy difference of ∼ 4.6% for CV2 (Welch’s t-test, t(20.945) = 5.217, p-value < 0.001) and ∼ 11.5% for CV1 (Welch’s t-test, t(22.205) = 6.817, p-value < 0.001); Fig. 4a, Table S6, Fig. S4). Using both SNPs and SVs as predictors has similar results as using only SVs.

**Fig. 4.**
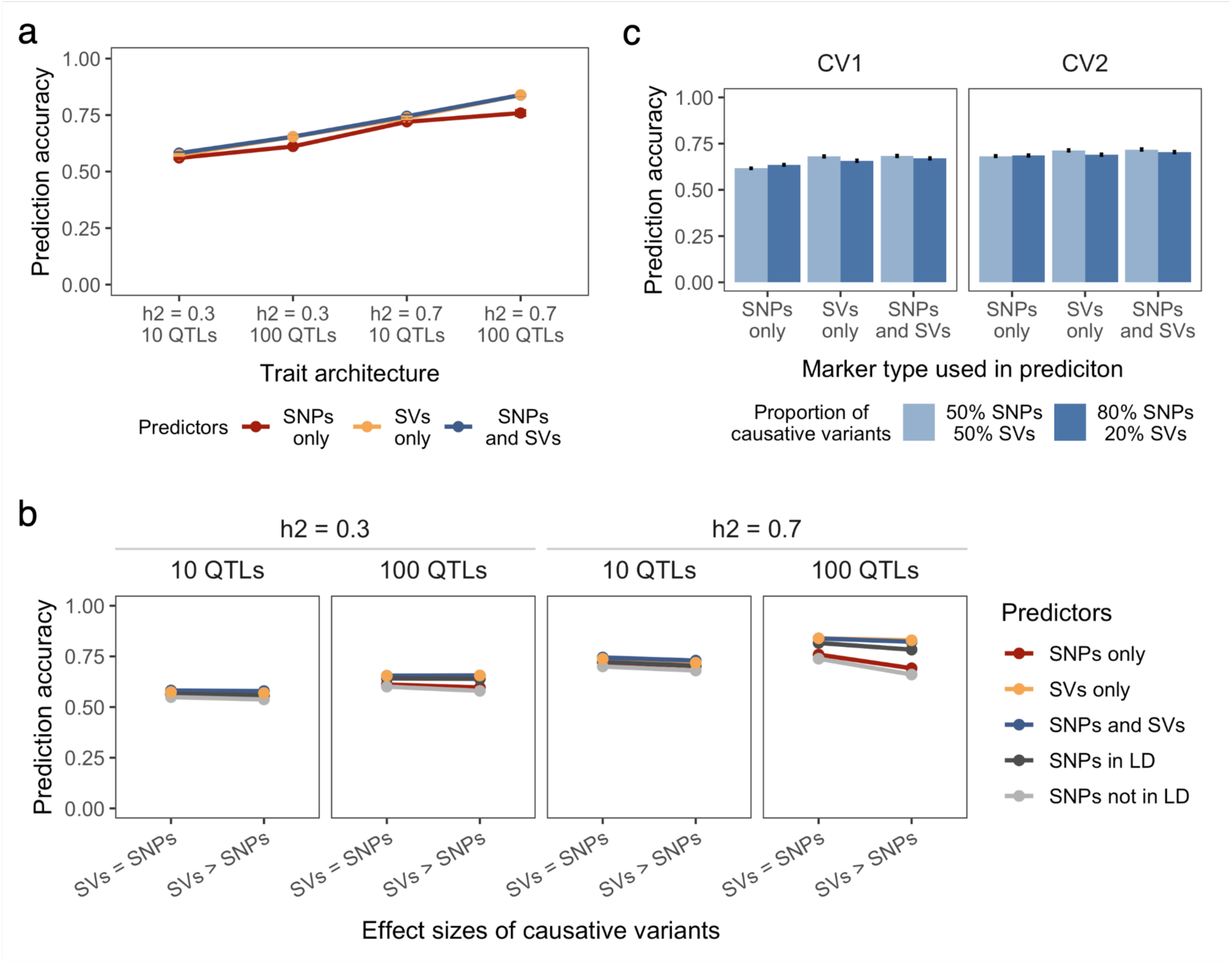
Genomic prediction accuracy of different marker types (SNPs and SVs, SNPs only, SVs only, SNPs in LD to SVs, or SNPs not in LD to SVs) from two cross-validation schemes (CV1 and CV2) for simulated traits where both SNPs and SVs were the causative variants. **a** Average prediction accuracy of marker types for each genetic architecture when the number and size of effects of both SNP and SV causative variant types are the same; **b** Changes in the average prediction accuracy after increasing the effect size of SVs relative SNPs. In both scenarios the proportion of SNP and SV causative variants was the same. In the SV > SNP effect size scenario the SV effect size was set to 5x that of the SNP effect size; c Average prediction difference between the two cross-validation schemes of different marker types when the proportion of causative variant types is the same or when most of causative variants are SNPs

We next wanted to investigate whether differences in effect sizes between SV and SNP causative variants would have an impact on prediction accuracy when using different marker types as predictors. Interestingly, we noticed that increasing the effect size of SVs relative to SNPs on trait variation only affects prediction accuracy in a scenario with high heritability and high number of QTLs (Fig. 4b, Table S6, Fig. S4). In this case, the accuracy of SNP predictors is lower (accuracy difference of ∼ 3.2% for CV2 (Welch’s t-test, t(32.81) = 2.381, p-value = 0.023) and ∼ 10.3% for CV1 (Welch’s t-test, t(33.87) = 3.728, p-value < 0.001)) while SV predictors have similar accuracy (accuracy difference of ∼ 0.1% (Welch’s t-test, t(35.515) = 0.181, p-value = 0.857) for CV2 and ∼ 1.8% for CV1 (Welch’s t-test, t(32.322) = 1.7843, p-value = 0.083)). We also observe a slight reduction in accuracy of using SNPs in LD to SVs as markers when the causal SVs have a larger effect than causal SNPs, likely due to the imperfect LD between SNPs and SVs, even for those in relatively high LD (Fig. 4b, Table S6, Fig. S4).

Finally, we tested the impact of predictor type on genomic prediction accuracy when there is an imbalance in the number of SNP and SV causative variants (Fig. S4). When there are more SNPs than SVs controlling trait variation (80% SNPs and 20% SVs), the benefit of using SVs as predictors reduces and its performance is more similar to using only SNPs (Fig. 4c, Table S6, Fig. S4). However, when predicting untested genotypes (i.e., CV1), SV predictors still perform better than SNP predictors (Fig. 4c).

Taken together, these results suggest that adding SVs to prediction models is beneficial when both SNPs and SVs are controlling highly heritable traits with many QTLs, regardless of the effect size difference or relative proportions of variants between causative SNPs and SVs.

### Linkage disequilibrium between causative variant and predictor, not predictor type, drives improvements in prediction accuracy

To further understand why different predictor types were able to achieve higher accuracies we explored the relationship between LD of causative variants and predictors in improving prediction accuracy. As expected, we observed that accuracy tends to increase as the number of predictors in LD to QTLs increases (Fig. 5a). Looking at the predictor in highest LD to a QTL in each simulation and prediction scenario and plotting the distribution of r^2^ values, demonstrated that most QTLs are being tagged in each scenario (Fig. 5b, Fig. S5). The high number of predictors and the big haplotype blocks of RILs increases the chance of having a causative variant tagged by a predictor. However, in a scenario where SNPs are the predictors and SVs are the QTLs (distribution highlighted in red in Fig. 5b), the number of predictors in very high LD to a QTL is considerably lower than other scenarios (e.g., when SVs are predictors), which may explain why SNP predictors do not perform very well when SVs are the causative variants.

**Fig. 5.**
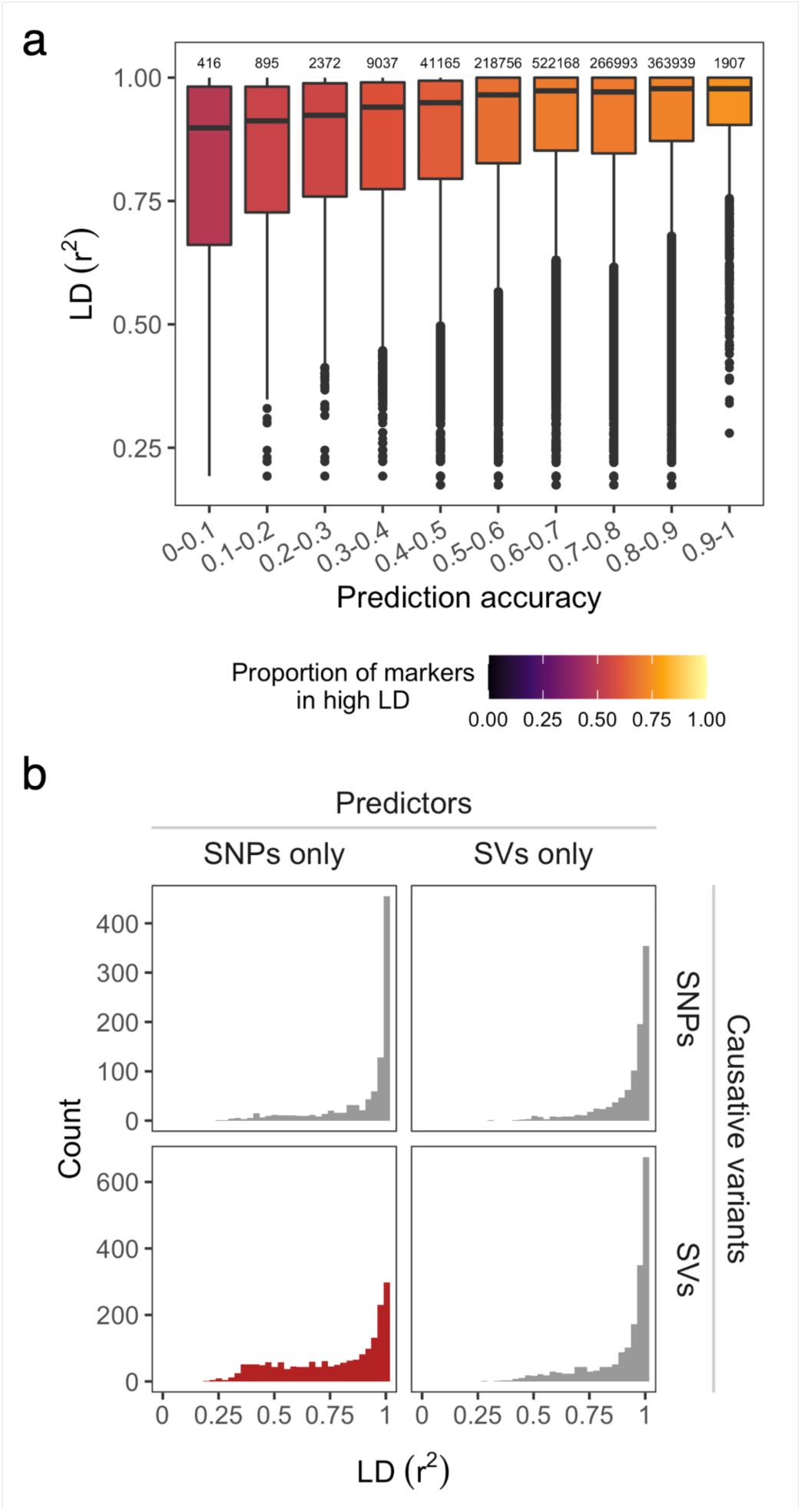
Relationship between genomic prediction accuracy and linkage disequilibrium (LD) of predictors and simulated QTLs. **a** Boxplot of pairwise r^2^ values between predictors and QTLs at different prediction accuracy bins. Numbers above each box represent the number of marker pairs with LD values; **b** Distribution of pairwise r^2^ values of SNP and SV predictors in highest LD to a QTL for all genetic architectures with either SNPs or SVs as causative variants

To further explore this observation in a more “real world” scenario, we downsampled the number of predictors to something more similar to what breeders use in breeding programs (i.e., 500 markers), and ran genomic prediction models with markers with different LD levels to a QTL (low, moderate or high). As expected, using only markers in high LD to a QTL resulted in the highest prediction accuracy regardless of marker type, followed by markers in moderate LD, and then markers in low LD e (Fig. 6, Table S7, Supplemental File 9). These results offer an explanation as to why SV markers outperform SNP markers in genomic prediction models when the causative variants are SVs and why SNPs in LD are able to provide comparable accuracies to actual SV predictors.

**Fig. 6.**
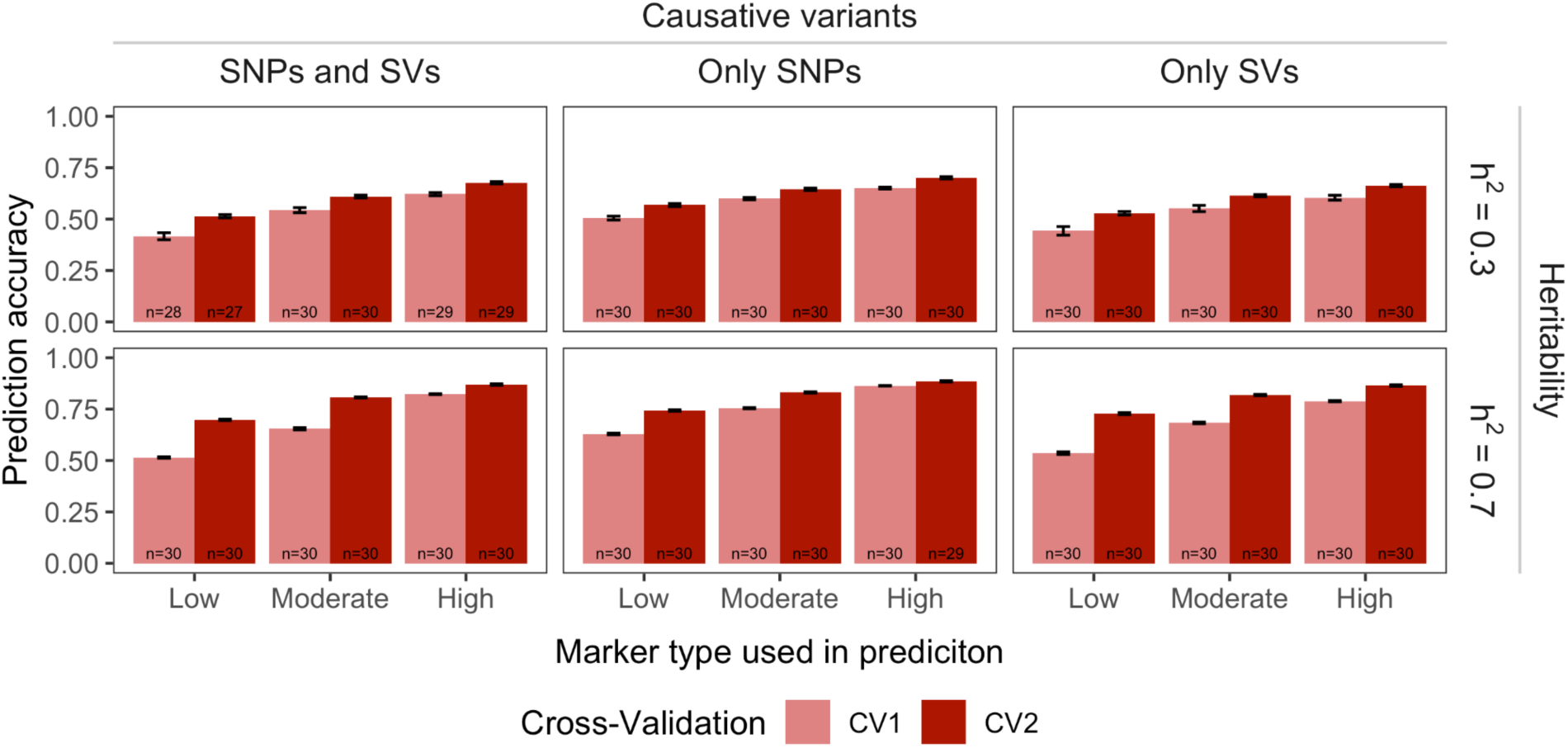
Genomic prediction accuracy from two cross-validation schemes (CV1 and CV2) of markers with low (r^2^ < 0.5), moderate (0.5 < r^2^ < 0.9) and high (r^2^ > 0.9) linkage disequilibrium (LD) to a QTL. Traits were simulated to have 100 QTLs, which could have been both SNPs and SVs (1:1 SNP-to-SV ratio), all SNPs, or all SVs. Numbers at the bottom of the bars represent how many scenarios the genomic prediction models successfully converged

## Discussion

In this study, we used a simulation approach to comprehensively evaluate the usefulness of structural variants in genomic prediction models as a complement to or in lieu of using SNP markers as predictors. One of the main reasons SNPs have been successfully used in genomic prediction models is their ubiquitous distribution across the genome, which maximizes the chance of having a SNP in close proximity and in linkage disequilibrium to a causal variant. However, previous studies showed that this is not always the case. For example, in a maize population of 103 lines, 20% of read-depth variants (a proxy for CNVs) could not be tagged by SNPs (Chia et al. 2012). More recently, a study showed that nearly 22% of polymorphic SVs in a diverse maize population of 521 lines were in low linkage disequilibrium with SNPs (Yang et al. 2019). Further, Windhausen et al. (2012) and Technow et al. (2014) showed that LD decays very fast even in closed breeding populations. We observed a similarly fast LD decay in our RIL population with ∼24% of segregating SVs in the multiparent population not being tagged by a SNP. These results suggest that SVs have the ability to bring additional information to genomic prediction models that are not already captured by SNPs even in a breeding population.

To investigate how genomic prediction models could benefit from the extra information that SVs could bring, we simulated traits with a range of different genetic architectures. Overall, traits with high heritability and high number of QTLs controlling trait variation yielded higher overall prediction accuracy than traits with low heritability and low number of QTLs regardless of the predictor type. The importance of heritability in prediction models has been shown previously (Daetwyler et al. 2010; Combs and Bernardo 2013; Guo et al. 2014), and the increased accuracy for higher heritabilities is expected due to the stronger influence of genetic effects over environmental effects on trait variability (Combs and Bernardo 2013). Although we also observed an increase in accuracy for traits with a high number of QTLs, this increase was not as evident as for changes in heritability. Others have shown that the number of QTLs had no to little effect on prediction accuracy (Daetwyler et al. 2010; Wientjes et al. 2015). Interestingly, Wientjes et al. (2015) showed that prediction accuracies were slightly higher for traits with 1000 QTLs compared to 100 QTLs when small effects were assigned to QTLs regardless of their allele frequency, which is a similar scenario to what we simulated in this study.

The main finding from our simulations with regards to predictor type was that the extent to which the extra information brought by SVs can be useful in prediction models depends on the genetic architecture of the trait. When traits were simulated to have high heritability and high number of QTLs the differences in prediction accuracy among different predictors became more evident, especially when SVs were the causative variants. For example, our findings suggest that if a low number of SVs control phenotypic variation and the heritability of the trait is low, then using SVs as markers for prediction models will not bring any additional information to the models. This architecture might explain, at least partially, why very modest or no improvement was observed in previous empirical studies using SVs in models (Lyra et al. 2018; Chen et al. 2021). Another possible explanation is that the SV markers used in these studies are not the causative variants or are not in high LD with the actual causative variants. Pérez-Enciso et al. (2015) showed in their simulations that prediction accuracy could increase by about 40% if all causal SNPs are included in a genomic prediction model, but it drops very quickly if such SNPs are removed, and non-causal variants are included. Similarly, van den Berg et al. (2016) also showed that genomic prediction reliability increases when markers closer to causative variants are included and concluded that markers should be carefully selected in prediction models. We also showed that selecting predictors in highest LD to causative variants, regardless of variant type, results in significant gains in prediction accuracy. Recently, Ramstein and Buckler (2021) showed that prioritizing genomic markers associated with variants most likely to impact trait variation can increase genomic prediction accuracy. Taken together these findings highlight the importance of understanding trait genetic architecture to maximize genomic prediction accuracy by carefully selecting predictors. As more association studies demonstrate how prevalent SVs are in genetic architectures (Yang et al. 2019; Alonge et al. 2020; Guo et al. 2020), breeders can make an informed decision about the use of SVs or SNPs in LD with SVs when there is information supporting SVs underlying their trait of interest.

Access to pangenomes that capture structural variation is now a reality for many crop species (Bayer et al. 2020; Della Coletta et al. 2021), and high quality SV data will become increasingly more available in the near future for many other crop species. Given the results described here, using SVs in prediction models in a breeding program can improve genomic prediction accuracy, but the added benefit of these markers may not yet be cost-effective to obtain at the scale of a commercial breeding program. For crop species already with large pan-genome information, such as maize, soybean, and tomato (Gao et al. 2019; Liu et al. 2020; Hufford et al. 2021), practical haplotype graphs (Franco et al. 2020; Jensen et al. 2020) may be a starting point to impute SV information from a representative set of individuals to breeding lines. As for actually genotyping the training and test populations, selecting SNP markers in LD to SVs has the potential to be used as a proxy to SVs without compromising accuracy and keeping genotyping costs low. Reaching near-perfect prediction accuracy is going to involve incorporating the maximum amount of non-redundant genetic and environmental information into the models. The results of this study are a step forward towards developing strategies that take advantage of all relevant information available in the plant genome by exploring the use of alternative marker types under varying genetic architectures.

## Acknowledgements

We thank DOW AgroScience (now Corteva Agriscience) for providing in-kind support through the custom Illumina Infinium 20k SNP chip. We thank the Minnesota Supercomputing Institute at the University of Minnesota (http://www.msi.umn.edu) for providing resources that contributed to the research results reported in this article.

## Statements and Declarations

### Authors’ contributions

Conceptualization: RDC, MAM, MOB, AEL, CNH; Methodology: RDC, SBF, PJM; Formal analysis and investigation: RDC, PJM; Writing - original draft preparation: RDC; Writing - review and editing: RDC, SBF, PJM, MAM, MOB, AEL, CNH; Funding acquisition and supervision: MAM, MOB, AEL, CNH. All authors read and approved the final manuscript.

### Availability of data and material

The datasets analyzed in this study are available in the Data Repository for U of M (DRUM) at https://hdl.handle.net/11299/250568 (genotypic data of maize parental lines and RILs (Della Coletta et al. 2023)) and https://conservancy.umn.edu/drum (Supplemental files 1-10 generated for this manuscript that includes structural variant calls of the maize parental lines, projected genotypic data for recombinant RILs, simulated trait values for each RIL with different genetic architectures, input data for genomic prediction models with different marker types, and genomic prediction accuracy for each combination of simulated genetic architecture and predictors.)

### Code availability

The codes/scripts used for data analysis are available in the Github repository https://github.com/HirschLabUMN/genomic_prediction_svs.

### Ethics approval

Not applicable

### Consent to participate

Not applicable

### Consent for publication

Not applicable

## Supplementary Information

**Fig. S1.**
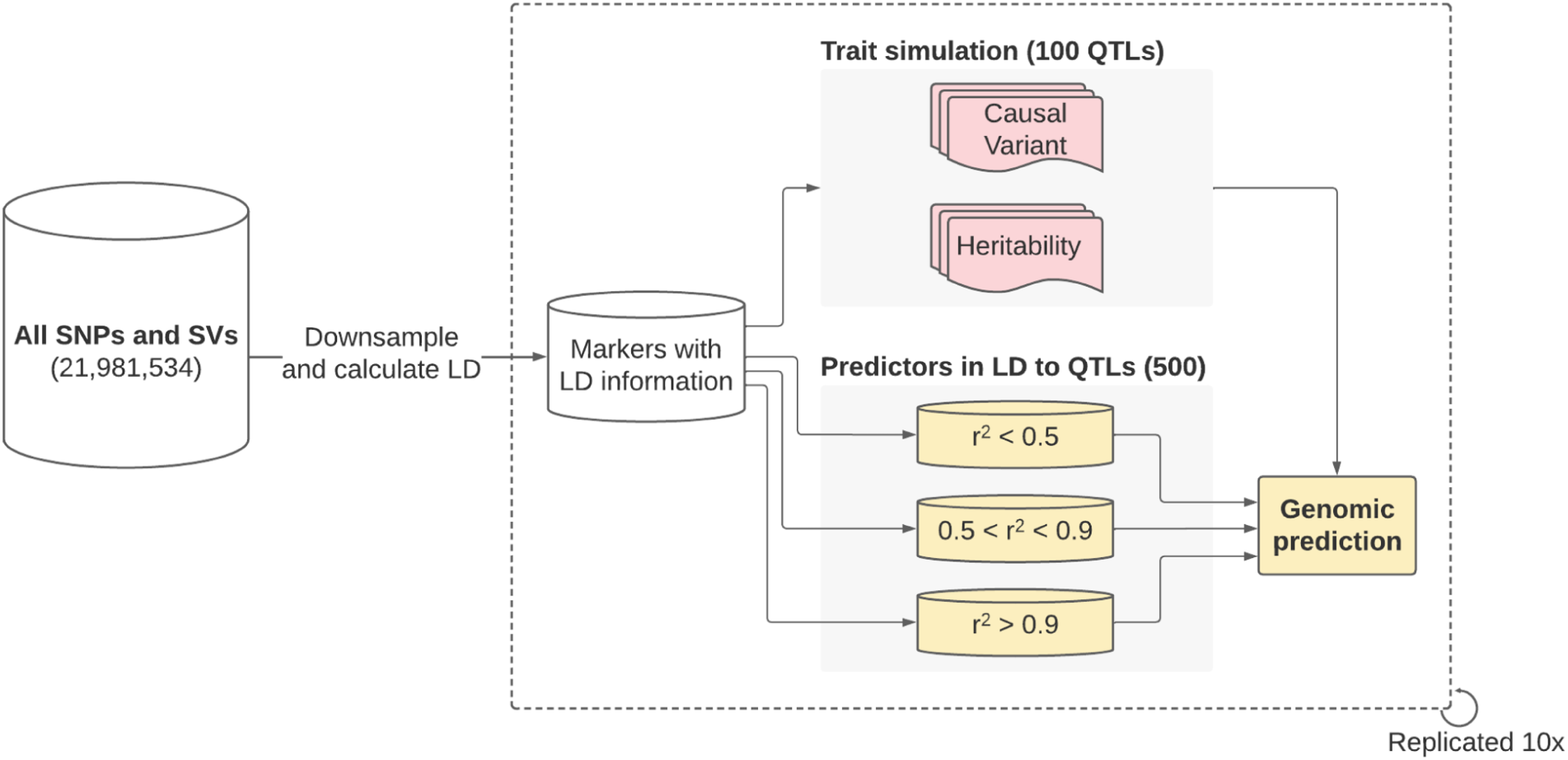
Methodology overview for simulating traits and testing performance of markers with different LD to a QTL in genomic prediction models

**Fig. S2.**
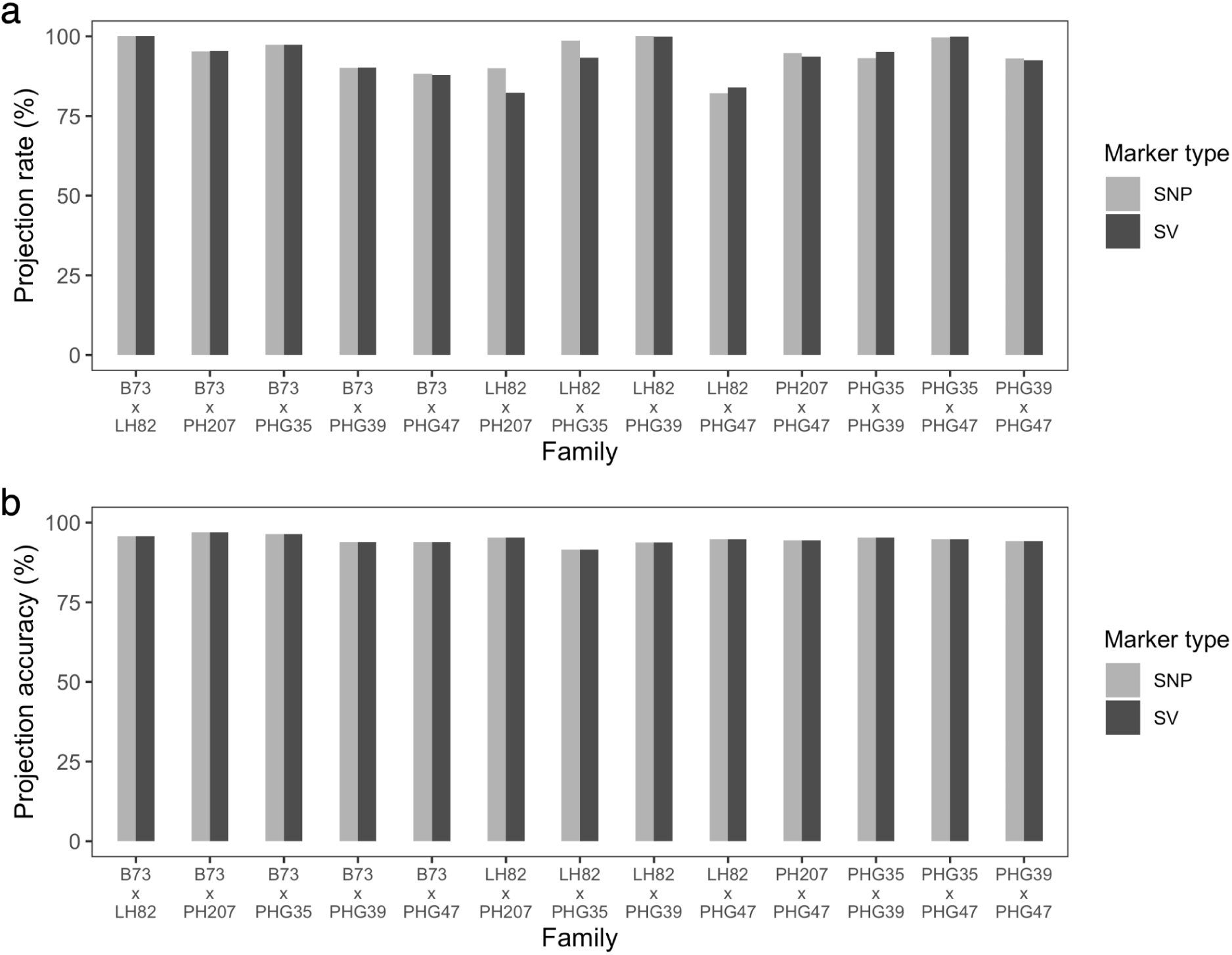
Projection rate (**a**) and accuracy (**b**) of SV and SNP markers from parents of 13 biparental populations to their respective recombinant inbred lines

**Fig. S3.**
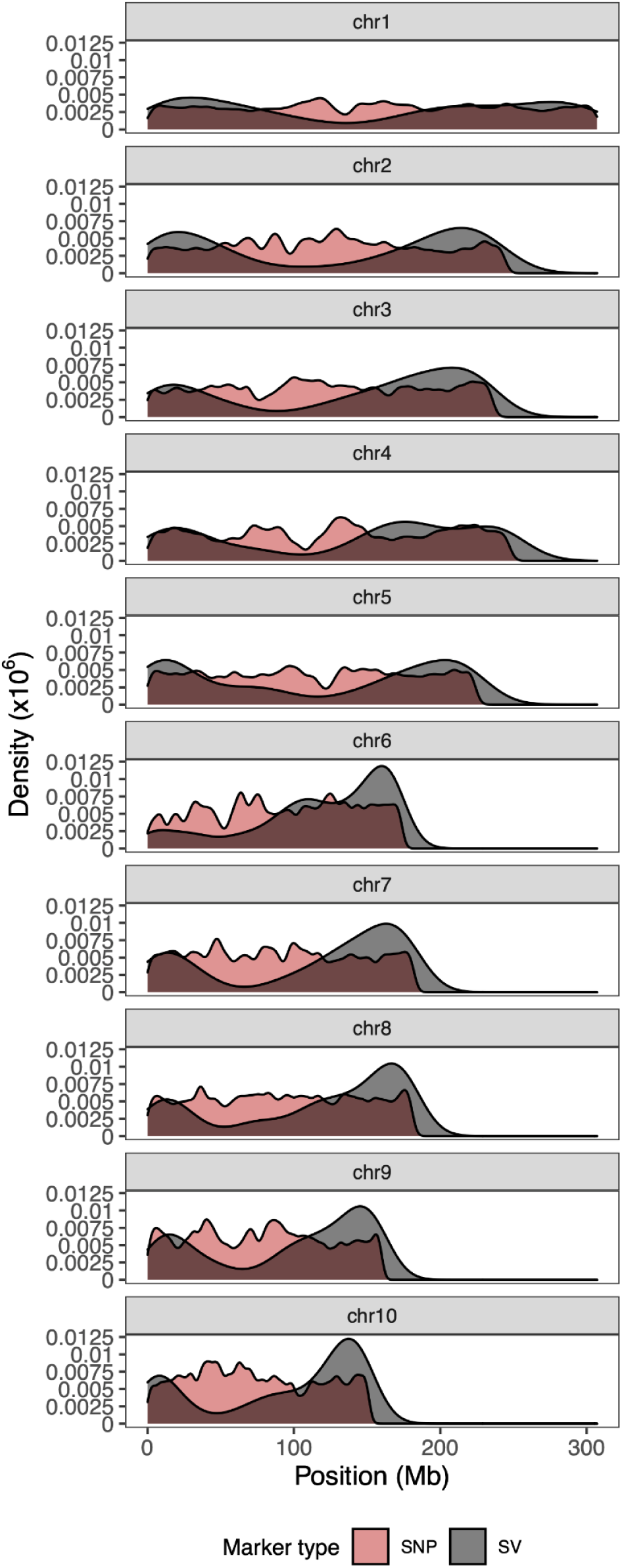
Density plots of all projected SNP and SV markers showing their distribution along the ten maize chromosomes

**Fig. S4.**
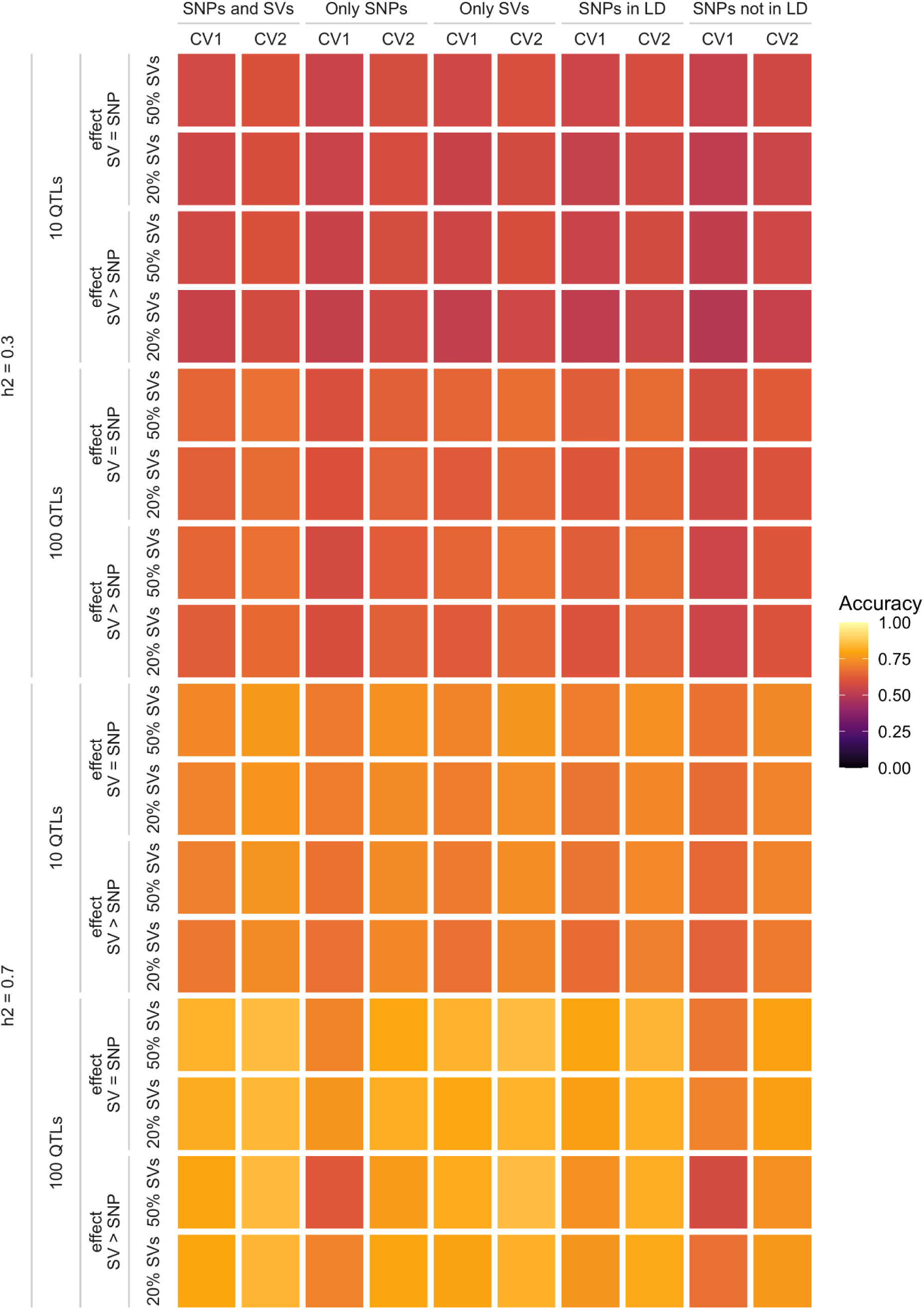
Heatmap summarizing genomic prediction accuracy from two cross-validation schemes (CV1 and CV2) of five marker types (SNPs only, SVs only, SNPs and SVs, SNPs in LD to SVs, and SNPs not in LD to SVs) used to predict phenotypic values of simulated traits controlled by both SNPs and SVs with the same or different effect sizes between SNP and SV causative variants. In the SV > SNP effect size scenario the SV effect size was set to 5x that of the SNP effect size

**Fig. S5.**
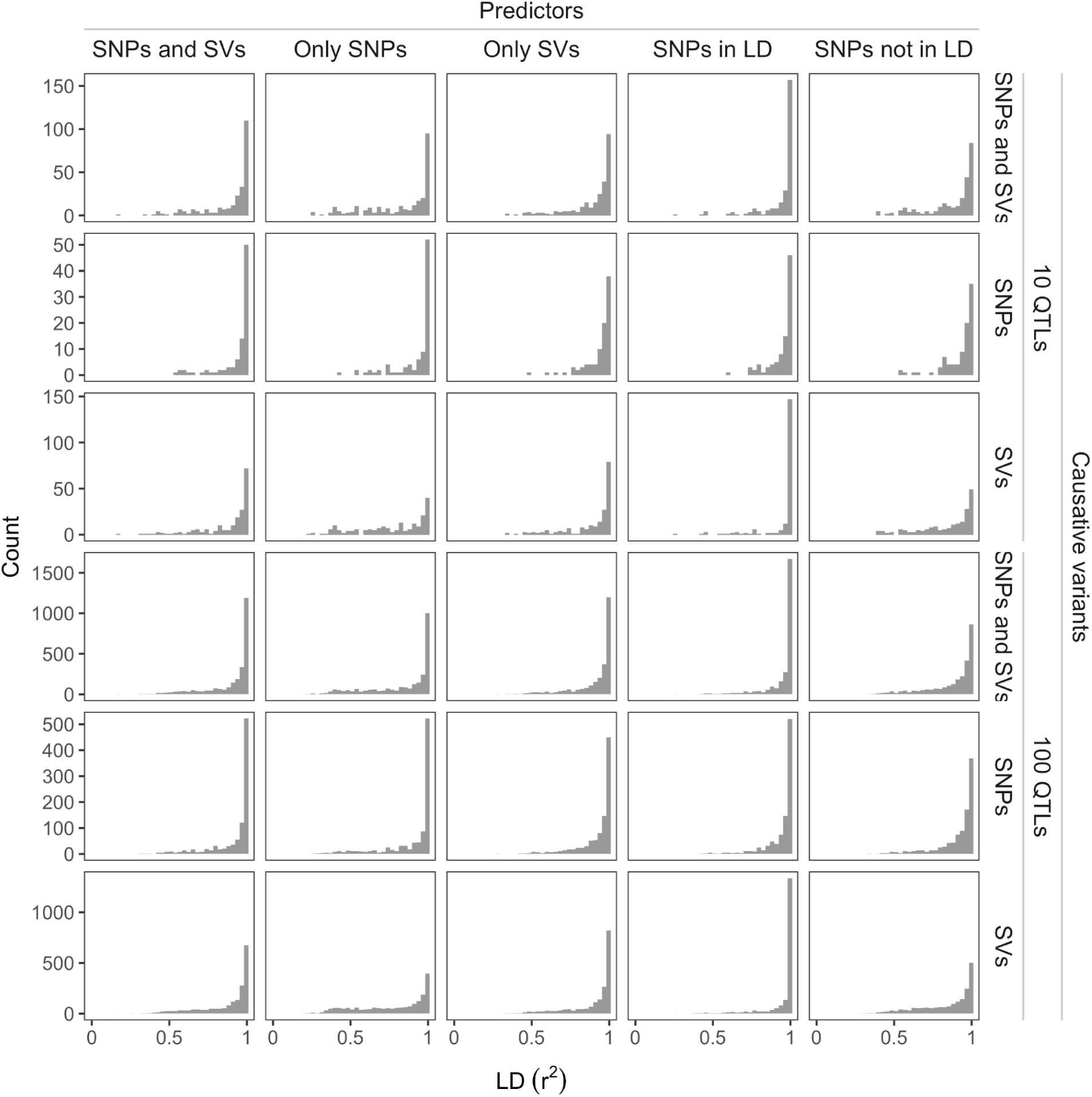
Distribution of pairwise r^2^ values of predictors in highest LD to a QTL for each genetic architecture

